# Social network structure scales with group size in a multi-species analysis

**DOI:** 10.1101/2023.08.28.555173

**Authors:** Brenna Marie Gagliardi, Nader St-Amant, Roslyn Dakin

**Author notes:** To whom correspondence should be addressed: Roslyn Dakin.

## Abstract

Social networks can shape the evolution and transmission of behaviours. Recent studies have characterized social networks within species, but we know relatively little about the drivers of variation in the structure of social networks across species. Here, we analyze a database of 631 social networks from more than 30 animal species to test how social network properties scale with group size. We examine three properties of social network topology: the uniformity of edge weights within the group, the selectivity of individuals for particular social partners, and the amount of heterogeneity among individuals in social behaviour. We show that most of the variation in these three network properties is due to differences between focal species and/or study methodologies, with only negligible differences between the three animal classes in our analysis (birds, mammals, and insects). Our analysis also indicates that group size is the key factor that determines the topology of animal social networks. In smaller groups, edge weights are distributed more uniformly, and individuals show greater selectivity regarding their social partners. These results suggest that there are general scaling rules governing the social networks of diverse animal species. In small groups, individuals form strong connections to a selective set of partners, whereas in large groups, weaker ties are common, and individuals are less likely to limit their interactions to specific partners.

**Significance Statement:** Sociality is observed throughout the animal kingdom. Animal societies also vary widely in group size, from a few individuals to groups with millions on individuals. How do animal social networks vary across this spectrum? We investigated this question using data compiled from recent bird, mammal, and insect studies. Surprisingly, we found broad overlap in the social network structures of these three distinct animal classes. We tested the effect of group size on social network structure, and found that in small groups, individuals form stronger social ties with preferred social partners; by contrast, animals in large groups have weaker and less selective social ties. These results suggest that diverse animal groups are governed by similar processes that determine the structure of their social networks.

## Introduction

Social structure, or the patterns of relationships between individuals in a population, plays an important role in many ecological and evolutionary processes (Croft et al. 2008; Kurvers et al. 2014). These can include disease transmission (Griffin and Nunn 2012; Sah et al. 2017; Sah et al. 2018), the transfer of information (Aplin et al. 2012; Farine and Whitehead 2015), and the evolution of cooperative behavior (Ohtsuki et al. 2006; Carter and Wilkinson 2015). The investigation of social structure has gained more attention over the last two decades as biologists have adopted social network analysis, a quantitative framework for investigating the properties of social groups (Croft et al. 2008; Krause et al. 2009; Webber and Vander Wal 2019). A key question is what processes determine the development and dynamics of social network structures (Barabasi and Albert 1999; Pinter-Wollman et al. 2014; Ilany and Akçay 2016; Dakin and Ryder 2020; Piefke et al. 2021). In a pioneering study, Lusseau (2003) found evidence of scale-free network properties in a small population of bottlenose dolphins. Scale-free networks, first described by Barabasi and Albert (1999), are characterized by social groups wherein a small subset of individuals are highly connected, central hubs. Despite the broad importance of social structure in behavioural evolution, there are relatively few studies that compare social structure across species and taxonomic groups (Shizuka and McDonald 2012; Sah et al. 2017; Sah et al. 2018; Hobson et al. 2021).

One key factor that may determine the properties of animal social networks is group size. Animal societies can vary widely in group size, from groups with a handful of individuals to those with millions. A previous study the social wasp *Ropalidia marginata* examined variable colony size within this species, and found that the wasp colonies exhibited predictable changes in social network topology as colony size increased (Naug 2009). Here, we use an analysis of over 600 animal social networks from a recent repository, the ASNR database (Sah et al. 2019), to describe how social network properties vary with group size among well-studied animal taxa.

The ASNR repository gathers animal social networks from previously published studies (Sah et al. 2019). Each of these networks consists of nodes, representing individuals, connected by edges, representing their pairwise social associations (Croft et al. 2008; Krause et al. 2009). We focused our analysis on weighted social networks, wherein edge weights are assigned as a dyad-level characteristic quantifying the strength of pairwise social ties (Croft et al. 2008; Sah et al. 2019). We analyzed entries in ASNR from bird, mammal, and insect studies, because these three taxa were relatively well-represented with multiple weighted networks that varied in group size. We focused our investigation here on three properties of the weighted social networks that are quantified at the group level: edge weight uniformity, social selectivity, and individual heterogeneity. Together, these three properties describe how individuals in a group distribute their interactions with others. Additionally, these parameters can be described for any social network, regardless of the valence of the edges, or whether edges were constructed based on specific interactions, spatial proximity events, or some combination thereof.

The first network property we examine in this study, edge weight uniformity, is a measure of the typical strength of social cohesion within an animal group. To facilitate comparison across studies, we quantified edge weight uniformity as the median standardized tie strength. High values on this metric represent networks with relatively uniform distributions of edge weights (i.e., more of the edges in the network approach the weight of the maximum observed in the group). By contrast, low values on this metric represent networks with more skew in edge weights (i.e., a greater proportion of ties that are much weaker than the group maximum). In previous studies within animal populations, stronger social ties have been shown to be more stable through time (Dakin and Ryder 2020). Tie strength has also been shown to predict the amount of resource and information transfer between dyads (Aplin et al. 2012; Carter and Wilkinson 2013). Based on the observation of scale-free network structures in an early dolphin study (Lusseau 2003), we predicted that edge weight uniformity would decline with group size in our multi-species analysis.

The next property we examine in this study is social selectivity, which we define as the extent to which individuals in a population concentrate their social interactions on a subset of their partners. If individuals in a group largely interact uniformly with a set of partners the network would have low selectivity; by contrast, a network wherein individuals interact primarily with a specific subset of their partners would have high selectivity. High selectivity can occur as a result of specific partner preferences, individual recognition, hierarchical roles and/or specialization within animal social groups (Croft et al. 2005; Gokcekus et al. 2021; Piefke et al. 2021; Shimoji and Dobata 2022). We expected that selectivity would increase with group size, but that this may be taxon specific as individual recognition mechanisms are expected to vary among animal groups.

The third property we consider is individual heterogeneity in social behaviour, i.e., the extent to which groupmates differ in their social behaviour. In general, among-individual differences in behaviour are ubiquitous across animal species and behavioural contexts (Bell et al. 2009). When considering social behaviour, individual heterogeneity can occur if individuals have well-defined differences in status, resources, or specialization (e.g., when there is a division or labour) (Dall et al. 2012; Aplin et al. 2015; Dakin and Ryder 2020). To quantify among-individual heterogeneity based on social network structure, we used each node’s standardized strength or weighted degree (its sum of edge weights), and then calculated the index of dispersion for these values within a given network. In a previous study of social wasp colonies, Naug (2009) found that there was a trend of increasing heterogeneity with increasing group size. As such, we expected heterogeneity to increase with group size in our multi-species analysis.

The networks used in our comparative analysis were also analyzed by Sah et al. in their work establishing how animal social network topology can predict disease transmission (Sah et al. 2018). Here, we focus on how social network properties scale with the number of individuals sampled in a particular animal group, and whether these relations differ across animal taxa. To investigate whether scaling relationships could arise from random social connections, we also simulated random networks that corresponded to the observed networks in group size and edge density, but had their edge weights distributed at random.

## Methods

### Network data and properties

We obtained animal social network data from the Animal Social Network Repository (ASNR) in September 2021 (Sah et al. 2019). We focused our analyses on weighted social networks from the three best-sampled taxonomic classes: birds (44 networks from 8 species or flock types), mammals (304 networks from 19 species) and insects (283 networks from 6 species). These three classes each had sufficient sample size to examine variation in network properties (Figure 1A). All data processing and analyses were performed in R 4.2.2 (R Core Team 2021). About 66% of the network samples were derived from repeated measures in two particular studies: 244 contact networks were recorded from six *Camponotus fellah* ant colonies (Mersch et al. 2013) and 190 contact networks were recorded from a population of *Microtus agrestis* voles (Davis et al. 2015). The remaining 217 networks in our analysis were derived from the other studies. As described in below, in all of our statistical analyses, we accounted for nonindependence of repeated measures from the same study.

**Figure 1.**
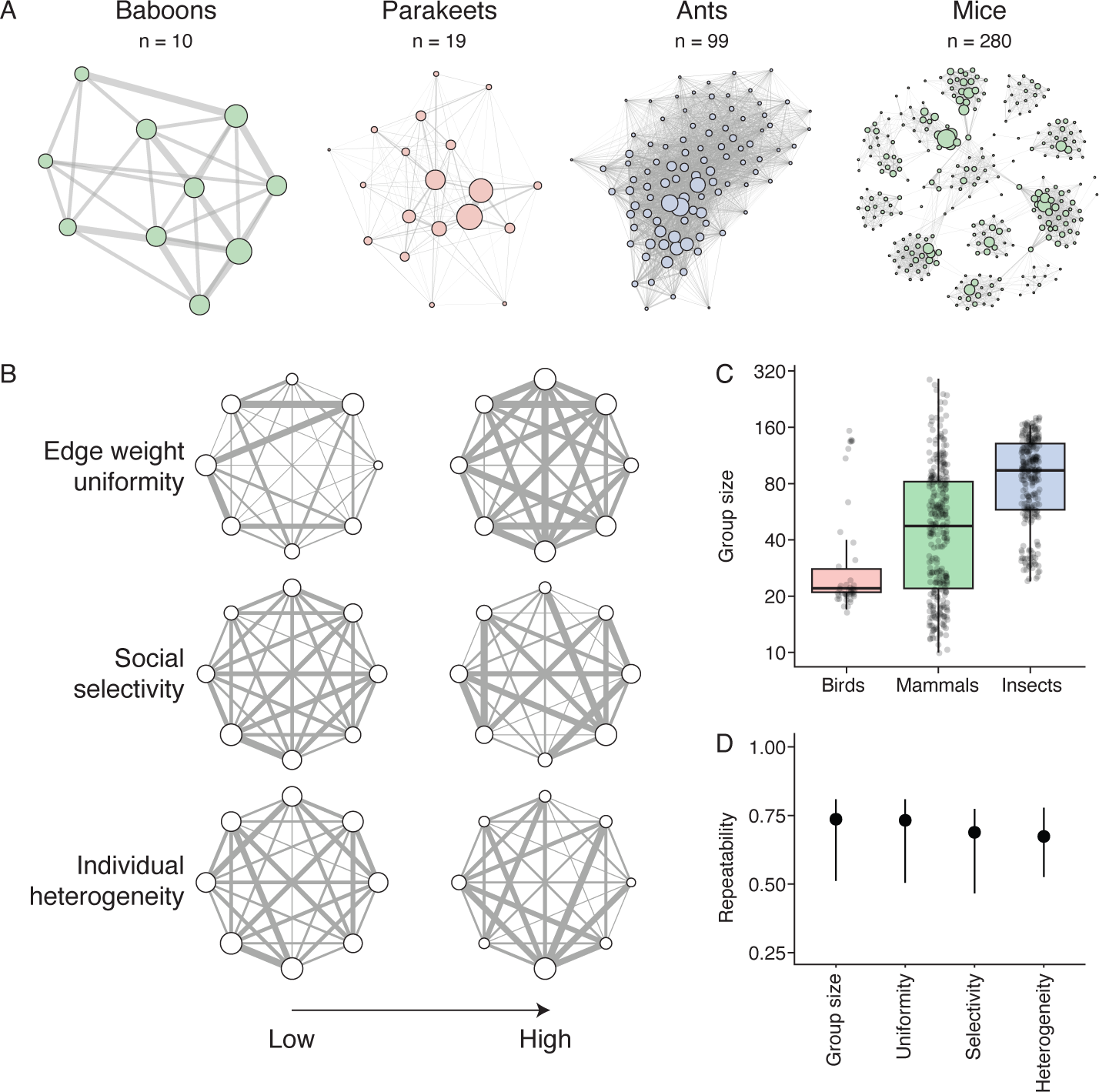
Animal social networks vary across studies. (A) Examples of four animal social networks from the ASNR database. Within each network, edge thicknesses are scaled to indicate variation in edge weight, and the size of each node represents its strength (i.e., the sum of its edge weights). Network layouts were determined with the Fruchterman-Reingold algorithm. (B) Theoretical examples of social networks with 8 individuals that vary in edge weight uniformity, social selectivity, and individual heterogeneity. In the top row, a network with high uniformity has fewer weak ties with low edge weight. In the middle row, a network with high selectivity has individuals that focus more of their social activity on a subset of their partners. On the bottom row, a more network with high heterogeneity has greater between-individual variation in social behaviour. (C) Group size for 631 animal social networks in our analysis. (D) Repeatability (R) estimates (± 95% CI) for the proportion of variation in network properties that can be attributed to study-level differences.

To allow comparison across networks, we first standardized the edge weights within each network to a maximum of 1, by dividing each edge weight by the maximum observed within the network. We assigned group size as the number of nodes (i.e., individuals) in the network. We used the package igraph 1.3.4 (Csárdi et al. 2022) to visualize examples for Figure 1A.

We calculated three network-level properties: edge weight uniformity, social selectivity, and individual heterogeneity (Figure 1B). We assigned edge weight uniformity by taking the median standardized edge weight of a network. Larger values of uniformity indicate that a greater proportion of the dyads in the network are relatively closely connected (Figure 1B). To quantify social selectivity for each network, we first computed node-level selectivity values by calculating the index of dispersion of each node’s non-zero edge weights. The index of dispersion is defined as the variance of a sample divided by its mean; hence, node-level selectivity increases if an individual’s edge weights are skewed toward a small subset of their partners. After computing node-level selectivity values, we took the average across all nodes in the network as the social selectivity of that network as a whole (Figure 1B). To quantify individual heterogeneity, we first calculated the index of dispersion of summed edge weights for each node (i.e., weighted degree values). Higher values of heterogeneity indicate that individuals vary more in their social activity (Figure 1B). By contrast, lower values of heterogeneity indicate that individuals in the network are more similar in terms of their social activity.

### Statistical analysis

Prior to further analysis, we natural log-transformed group size, uniformity, selectivity, and heterogeneity values, because they are strongly right-skewed. We verified that residuals met assumptions of all further statistical analyses.

We used repeatability analyses to examine how network properties differed among the major taxonomic groups and focal species/study methodologies. Although our sample size of social networks from the ASNR database was large (n = 631), it represents only 37 studies of 33 different focal species or flock types. Thus, species differences were confounded with study-level differences in methodology in the dataset. Given this confound, we refer to study-level differences (rather than focal species-level differences). To estimate taxon and study-level differences, we used the package rptR 0.9.22 (Stoffel et al. 2017) and entered taxon and study as random effects, running 1,000 parametric bootstrap iterations for each social network property.

To examine how social network properties scale with group size, we fit a series of mixed-effects regression models for uniformity, selectivity, and heterogeneity using the packages lme4 1.1-31 (Bates et al. 2015) and lmerTest 3.1-3 (Kuznetsova et al. 2017). All models had study as a random effect to account for nonindependence of repeated measures from the same study (and focal species). For each network property, we compared the following candidate models: (i) a model with taxonomic class as the only fixed effect; (ii) group size + taxonomic class; and (iii) group size * taxonomic class (i.e., allowing the scaling relationship to differ across taxa). We used AIC in the MuMIn package 1.46.0 (Bartoń 2022) to compare models, and visreg 2.7.0 (Breheny and Burchett 2020) to visualize model fits. We used the likelihood ratio test to evaluate the statistical significance of interaction terms. A small number of the weighted social networks in the ASNR database were based on mixed-species bird flocks (n = 7 out of 631 in our analysis). We verified that our conclusions did not change if these mixed-species samples were omitted. We also verified that conclusions were unchanged in the analysis of social selectivity if we omitted one mammal network that had an extreme low value for selectivity.

### Random networks

We assume that all animal social networks are governed by a combination of selective behaviours (directed at particular individuals), constraints (e.g., owing to the physical environment), and random chance. To determine what type of scaling relationships would be expected for social networks governed solely by chance, we simulated and analyzed a series of random networks. For each of the observed animal social network in our multi-species analysis, we simulated 100 random networks that had the same group size (N) and edge density but had edge weight units distributed among those edges at random. Our approach to defining edge weights was based on an initial assumption that individuals have some average frequency of interactions or contacts with others, such that the total number of contacts will scale with group size. We chose 3N as the number of edge weight units to distribute, because it represented a balance between too few units (which would mean that the networks would approach binary or unweighted networks) and a very large number of observations (which would mean that the random edge weight variation would approach a normal distribution). Thus, for a study that had a group size of 20 animals with 45 edges, each simulated random network also had 45 edges that would be placed at random, and the weights for those 45 edges would be determined by a random distribution of 60 edge weight units (60 = 3 x 20). After generating each simulated random network, we then standardized its edge weights to a maximum of 1, as was done with the observed animal social networks. Then, we calculated the same three properties of social network structure, defined above. For each observed network, we determined the average characteristics of 100 simulated random networks. We used paired t-tests to compare the characteristics of observed animal social networks with those of simulated random networks. We also repeated the scaling analysis on the simulated network properties to visualize the scaling relations expected in simple random networks.

## Results

Animal social network properties vary widely across studies, with about R = 67-74% of the variation in group size, uniformity, selectivity, and heterogeneity attributed to differences between studies (Figure 1D; all p-values < 0.0001). By contrast, when accounting for study-level differences, taxonomic class explained virtually none of the variation in any of these four social network properties (all R estimates = 0% and p-values > 0.99). Real animal social networks also have lower edge weight uniformity, greater social selectivity, and greater individual heterogeneity as compared to simulated random groups (paired t-tests, all p < 0.0001; Figure 2). Edge weight uniformity scales negatively with group size in a consistent manner across the social networks of birds, mammals, and insects. All else being equal, larger animal groups have less uniformity (and hence weaker social ties) as compared to smaller groups (Figure 2A, Tables 1-2). There was some weak support for a taxon-specific slope for uniformity and group size (Akaike weight 0.29 for the interaction model), but a test of the group size * taxon interaction in that model failed to reach statistical significance (likelihood ratio test, p = 0.10).

**Figure 2.**
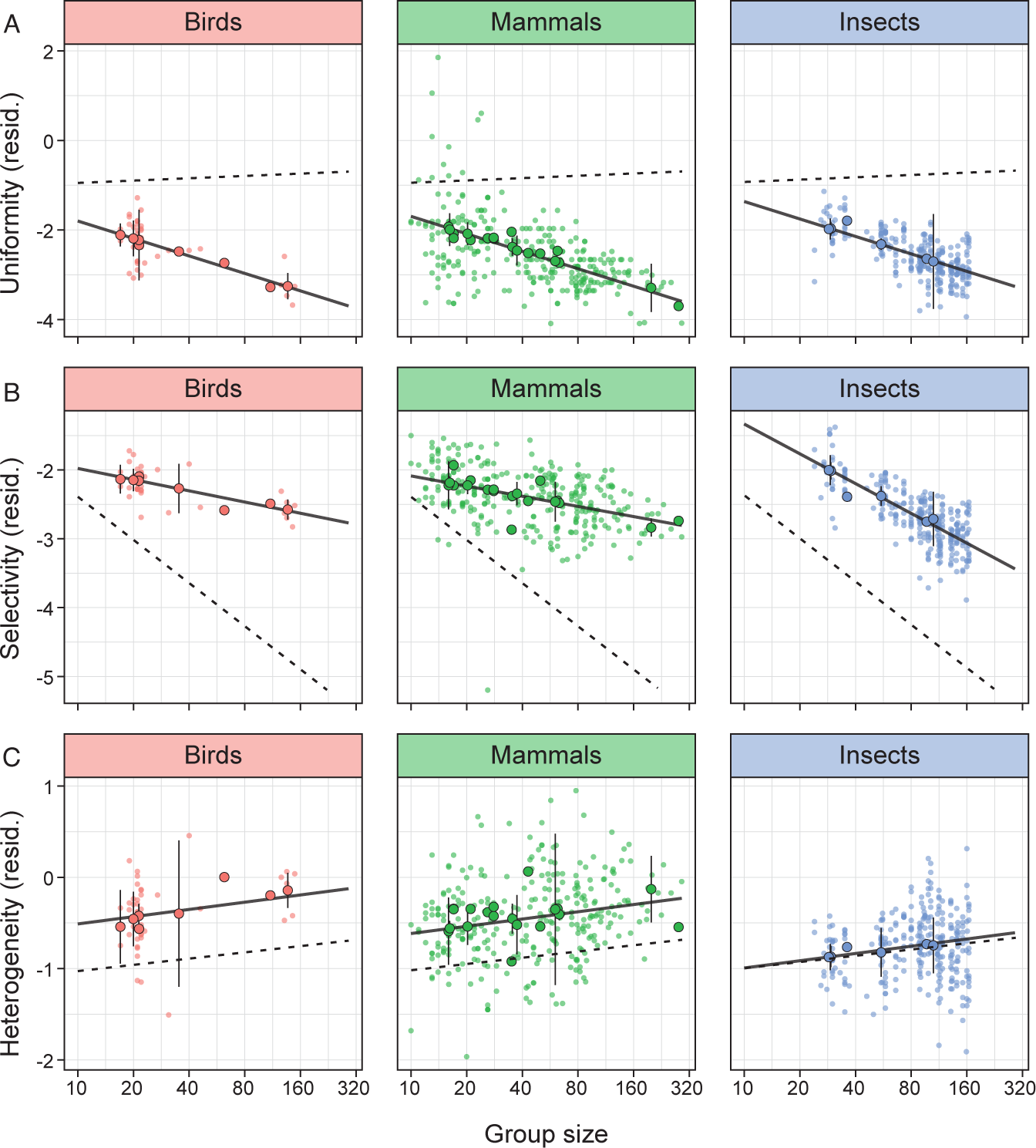
The properties of social networks scale with group size. (A) Edge weight uniformity, (B) social selectivity, and (C) individual heterogeneity in relation to group size. Each panel shows a partial residual plot from the best-supported mixed-effects model in Table 2 analyzing natural-log transformed metrics, with partial residuals on the y-axis and model predictions shown as black lines (n = 631 networks from 37 studies). The larger points with black outlines represent species-averages (± 95% CI). Dashed lines show expectations based on simulated networks where edge weight is distributed randomly.

**Table 1.**
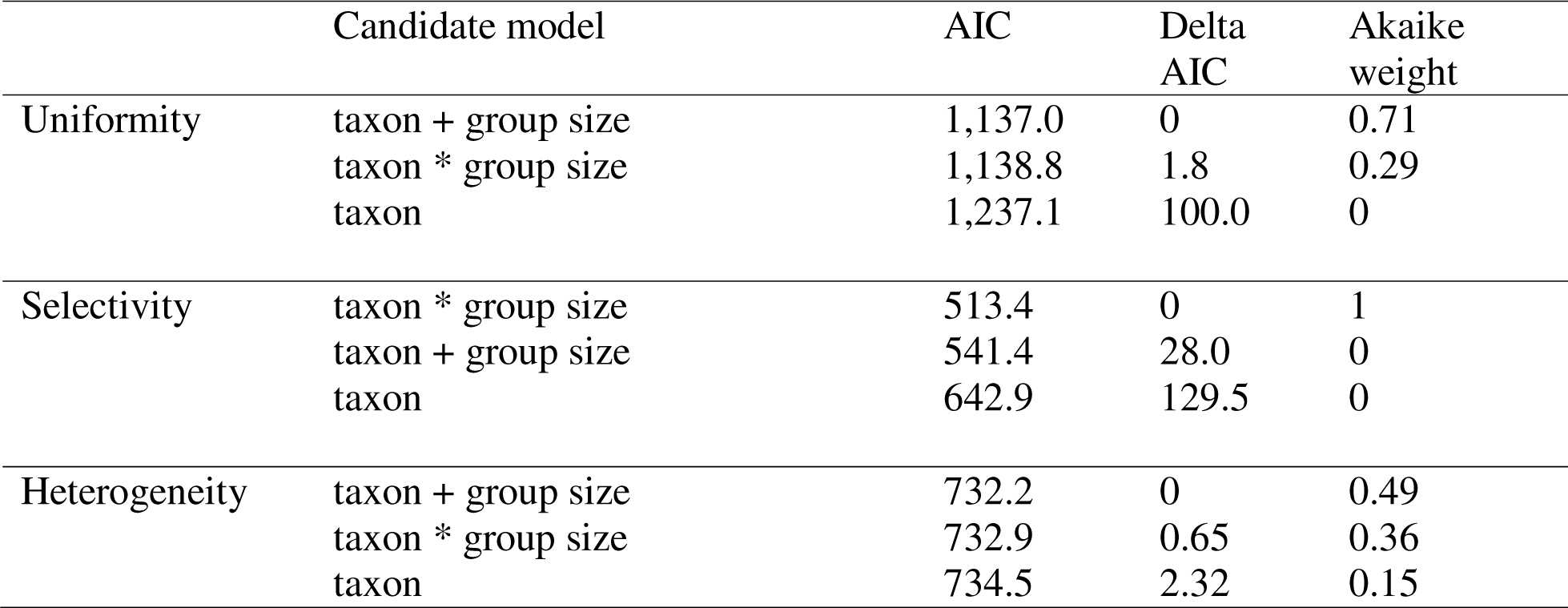
Candidate models for the scaling of social network properties with group size.

**Table 2.** Best-supported models of social network properties. All network properties and group size were natural log-transformed prior to analysis. Taxon is a categorical predictor with three levels (birds, mammals, and insects), and “Birds” is the reference category for estimates reported within this table. The sample size for these analyses is n = 631 networks from 37 studies.

|  | Predictor | Estimate | SE | t | p-value |
| --- | --- | --- | --- | --- | --- |
| Uniformity | intercept | -0.50 | 0.42 | -1.20 | 0.23 |
|  | taxon (Mammals) | 0.10 | 0.44 | 0.23 | 0.82 |
|  | taxon (Insects) | 0.44 | 0.58 | 0.75 | 0.46 |
|  | group size | -0.56 | 0.05 | -10.79 | < 0.0001 |
| Selectivity | intercept | -1.44 | 0.93 | -1.54 | 0.13 |
|  | taxon (Mammals) | -0.16 | 0.95 | -0.17 | 0.86 |
|  | taxon (Insects) | 1.54 | 0.99 | 1.56 | 0.13 |
|  | group size | -0.23 | 0.25 | -0.93 | 0.36 |
|  | group size : Mammals | 0.02 | 0.25 | 0.09 | 0.93 |
|  | group size : Insects | -0.39 | 0.26 | -1.51 | 0.14 |
| Heterogeneity | intercept | -0.77 | 0.24 | -3.18 | 0.002 |
|  | taxon (Mammals) | -0.111 | 0.24 | -0.43 | 0.67 |
|  | taxon (Insects) | -0.48 | 0.31 | -1.54 | 0.13 |
|  | group size | 0.11 | 0.04 | 3.04 | 0.002 |

Social selectivity also scales negatively with animal group size: the larger the group size, the less biased individuals are toward particular social partners (Figure 2B, Tables 1-2). The slope of the selectivity-group size relationship also differs across taxa (likelihood ratio test, p < 0.0001), with strong evidence in favour of the model with a group size * taxon interaction (Akaike weight = 1). Examining these slopes, we see that insects exhibit the steepest decline in social selectivity as group size increases (Figure 2B).

Individual heterogeneity tends to increase slightly with group size: larger groups tend to have greater differences among individual animals in the network (Figure 2C). We found some evidence that the slope of this relationship may differ across taxa (likelihood ratio test, p = 0.02), but the support for this was fairly weak and the best-supported model of heterogeneity was the model without an interaction (Figure 2C, Tables 1-2).

## Discussion

Here, we use a comparative multi-species approach to study how social network structure varies across birds, mammals, and insects. We focused our analysis on general properties that describe the distribution of social connections within groups. Surprisingly, we found broad overlap between the three taxonomic groups (i.e., there is as much variation within each taxonomic class as there is in the entire distribution of animal social networks sampled here). Most of the variation in social network properties is due to differences between studies (Figure 1D). This is consistent with the recent findings of Hobson et al. 2021, who found in their comparative analysis of dominance hierarchies that emergent features of dominance networks were not phylogenetically constrained.

Uniformity and selectivity characterize the relative strength of ties, and the extent to which individuals focus their interactions on specific partners. Our results revealed that these two traits both follow a similar scaling rule across taxonomic orders: all else being equal, edge weight uniformity and selectivity decline in a log-linear relationship with group size (Figure 2A-B). What behavioural processes drive these general scaling patterns? When social interactions are unconstrained, individuals are expected to accumulate more weak ties; and more weak ties would decrease our measure of average uniformity (or median standardized connection strength within a network). Similarly, when individuals are less able to limit their interactions to particular social partners, network selectivity will also decline. We propose that the scaling relations in Figure 2A-B derive from a mechanism whereby individuals in small groups tend to form strong connections with preferred partners, whereas those in large groups are either less likely (i.e., due to chance), or less able to limit their interactions to specific partners, and consequently make more frequent contacts with weak ties.

Differences in selectivity may also be attributed to differences in individual recognition and social cognitive abilities (Tibbetts and Dale 2007; Emery and Clayton 2009). If a group scores highly on social selectivity, individuals may either be constrained in who they interact with (e.g., spatially), or they may be able to recognize and return to specific partners repeatedly. Animals interactions are not solely dictated by chance, but rather animals choose who they interact with based on prior interactions and their level of social cognition (Hobson et al. 2021). As an example, Casey et al. (2015) found that elephant seals associate the cry of another with the outcome of a dominance contest. In general, larger groups may offer less opportunity for individuals to form connections with known partners. This could form an interesting avenue for further research on the relationships between selectivity, social cognition, and group size, as partner recognition is a key prerequisite in models of the evolution of cooperation based on reciprocity and reputation.

One difference between taxa in our analysis is that insects were found to have a steeper decline in selectivity as group size increases, as compared to mammals and birds (Figure 2B; this is also shown by the significant interaction in Table 2). Notably, the lowest values of selectivity, and many of the largest group sizes, are also found among very large eusocial insect groups. We are cautious in interpreting this taxon difference, given that the scaling relationship for insects in our analysis was also dominated by a single study (Mersch et al. 2013). On the other hand, ants account for a third of all insect biomass and far surpass that of land vertebrates (Wilson and Hölldobler 2005), making it reasonable to assume that eusocial insect colonies comprise the majority of large animal social groups worldwide. While eusocial insects have specialized roles, division of labour, and sophisticated communication abilities, the benefits of social interactions in eusocial insect taxa are often driven by inclusive fitness and high levels of kinship shared by the workers in these systems (Hamilton 1964). Hence, unlike systems where the benefits and costs are driven by reciprocity, reputation, and/or differences in kinship, we generally do not expect eusocial insects to seek out specific partnerships within their colonies (Crowley et al. 1996; Keller 1997; Tumulty et al. 2021).

The third property in our study, individual heterogeneity, captures the extent to which individuals in a group differ in their social behaviour. A group with high heterogeneity is one wherein some individuals are much more central than others; when a group scores low on heterogeneity, individuals are more similar in their level of social interactivity. Our analysis indicates that heterogeneity generally increases with group size (Figure 3C), which is in line with the previous findings of Naug (2009). In larger groups, there may be may be more opportunity for among-individual differentiation based on social status, differential resource access, or specialization (Fewell 2003; Naug 2009). Naug (2009) showed that social wasps in smaller groups had a tendency to take more generalist approaches, whereas in larger groups, individuals were more likely to specialize. In smaller groups, there may also be greater opportunity for coordination and phenotype matching, which would attenuate heterogeneity. It is important to note that while the slope of the heterogeneity-group size relationship was statistically significant in our multi-species analysis, it was very weak. Hence, much of the variation in heterogeneity for the diverse animal networks here was not explained by group size (Figure 3C, Table 2).

We also compared our results to simulated networks where edge weight variation was distributed randomly, represented by the dotted lines in Figure 2. We found that the real animal social networks are less uniform, more selective, and have greater among-individual heterogeneity than purely random networks. The decline in uniformity with group size found in real animal social networks is at odds with the expectation for random networks. By contrast, in random networks, selectivity is expected to decline with group size, and heterogeneity to increase, similar to the direction of change observed in actual animal groups. The social networks of birds and mammals exhibited milder selectivity declines with group size as compared to random networks, as observed by the comparison of slopes within Figure 2. Interactions in these bird and mammal groups are often governed by social information use, reciprocity, and/or spatial constraints, generating divergence from the random expectation (Dall et al. 2012; Aplin et al. 2015; Dakin and Ryder 2020). When examining the results for insects, we found a similar heterogeneity-group size relationship for real insect networks and random networks. Although it is well known that the social networks of eusocial insects are not purely random (Fewell 2003), chance may play a much larger role in their social interactions as they are less selective than other taxa (Keller 1997), bringing insects closer to the null expectation for the properties we analyzed.

Our study is just a first step into the exploration of cross-taxon trends in social structure, and there are many limitations that come with this. The social network parameters we chose are not the only ones possible. Sah et. al (2018) included a multitude of social parameters of relevance in their study on disease transmission in different social networks such as fragmentation, clustering, and network diameter, among others. We would expect that other social network properties would scale with group size across species. The ASNR database (Sah et al. 2019) which we used to perform this study is also dominated by a small number of species representing a large amount of the datapoints. It is possible that more studies on a wider variety of species may reveal additional features to these scaling trends. However, a final important note is that our present analysis cannot distinguish between variation caused by species and study methodology differences. With the rising number of social network studies being done as well as open data becoming more common, comparative cross-species studies such as this one are increasingly possible (Webber and Vander Wal 2019). There is a clear need for greater standardization in social network analysis practices, in order to open up these new research avenues on the mechanisms that broadly govern animal social structure (Webber and Vander Wal 2019).

## DATA ACCESSIBILITY

All data and R scripts are available at: https://figshare.com/s/9e80a039136bc236e5fb

## AUTHOR CONTRIBUTIONS

RD and BMG designed the study; RD and BMG analyzed the data; all authors wrote and edited the manuscript.

## ACKNOWLEDGEMENTS

We thank Pratha Sah, José David Méndez, and Shweta Bansal for developing the ASNR database, as well as all authors who provided their data for use in the database.

